# Augmenting visual errors or variability does not enhance motor learning in remote web application tasks

**DOI:** 10.1101/2024.05.10.593506

**Authors:** Nobuyasu Nakano, Akihiko Murai

## Abstract

Laboratory experiments employing robotic manipulandum are far from achieving their goal of helping people improve their motor learning. Remote experiments using web applications are an effective tool for bridging the gap between robotic manipulandum experiments in the laboratory and general motor tasks outside. However, the influence of interventions that increase error or variability in remote motor tasks on motor learning has not yet been determined. In this study, we aimed to elucidate the effects of interventions that visually increase errors and variability in remote experiments using web applications. In particular, 48 people participated in a web-based study on the cursor-manipulation of motor tasks using laptops. Three motor tasks (visuomotor-rotation reaching, virtual curling, and virtual ball-throwing tasks) were conducted, and each task consisted of 120 trials a day conducted for three days in this study. For each task, no intervention was provided on Day 1 and the intervention to augment motor error or variability was provided on Days 2 and 3. Differences between the groups in post-intervention test trials were examined using statistical analyses. Contrary to our expectations, the interventions of error-augmentation did not exhibit positive effects in Experiments 1 and 2, which could be attributed to a lack of haptic and proprioceptive information or inaccuracies in movement kinematics. In addition, the interventions of variability-augmentation did not exhibit positive effects in Experiment 3, which could be attributed to the complex dynamics in the relationship between perceived body movements and motor outcomes. Further research is required to identify the differences between the conditions when the interventions are effective or ineffective. Moreover, interventions must be developed to further improve general motor skills.

## 1 INTRODUCTION

In many sports and other scenarios, performers seek to minimize motor errors and variability. For example, in a motor task directed toward a target goal at a specific position, such as basketball shooting, motor errors and variability are directly related to performance. In general, learning a novel motor task reduces motor errors and variability in goal-directed movements during the learning process. However, in most cases, the improvement generally reaches a plateau early in the learning process and stops, such that only a few people attain elite performance, as demonstrated by top athletes. Thus, effective interventions are required to enable higher performance than that achieved via natural exercise. For example, when learning piano keystroke movements, the ability to discriminate the strengths of piano strokes improved in well-trained pianists who practiced with haptic devices (Hirano et al., 2020). The desired intervention is a system that is less dependent on the abilities of performers or coaches, and that can be used for multiple people.

To build a system that is not substantially dependent on individual ability, visual or dynamic (haptic) intervention using a computer or robotic device is suitable, as it eliminates the need for human verbal or physical instruction by a person (Heuer and Luettgen, 2015). In recent years, interventions that visually and dynamically increase motor errors and variability during learning have been reported to have a positive effect on motor learning (Sternad, 2018). In a force field reaching task, the amount and speed of motor adaptation were increased when learning was performed with visual interventions by displaying hand feedback at a virtual position that augmented the error of the hand positions along a straight-line trajectory to a target (Wei et al., 2005; Sharp et al., 2011). In a leg-step-like task using a robotic device, increased knee muscle activation and decreased leg tracking error were observed when learning was performed with haptic interverntions by forcing knee movements away from the correct position (Marchal-Crespo et al., 2014a,b). In addition, the effects of the interventions with visual and haptic error augmentation were compared in pinball-simulator (Milot et al., 2010; Bouchard et al., 2015) and gait-pattern modification tasks (Marchal-Crespo et al., 2019). Additionally, random noise that implicitly increases motor variability can drive motor learning toward decreasing motor variability, resulting in the acquisition of different movement patterns or strategies in a three-joint reaching (Mehler et al., 2017) and hand posture control tasks (Thorp et al., 2017).

Although numerous reports have revealed positive effects of the interventions with error or variability augmentation on motor learning, as discussed above, several studies have revealed negative effects. In a rowing task, in which participants are required to reproduce an ideal stroke, training with an intervention that visually amplifies the oar angle error has no benefit on stroke accuracy (Gerig et al., 2019). In a reaching task, where sound feedback is provided depending on success or failure (Therrien et al., 2018) and in a virtual shuffleboard task, where the disk speed was determined using the speeds of both hands (Cardis et al., 2018), random noise, which implicitly increases motor variability, impairs motor learning. These interventions may have negative effects; therefore, to develop interventions that are useful for actual motor learning, the effectiveness of the interventions must be tested by conducting experiments under the conditions similar to those of the target scenario.

Most conventional research, as described previously, employs robotic manipulandum developed for a specific purpose in the laboratory. Although such laboratory experiments are advantageous in that they strictly control subjects and accurately measure human movement, a large gap exists between these experiments and the goal of helping the public improve their motor learning. Within this context, the recent COVID-19 pandemic has increased the importance of non-face-to-face experiments, which has led to an increase in the number of remote experiments. Task programs for online reaching motor tasks (Tsay et al., 2020) and remote virtual reality (VR) motor tasks (Cesanek et al., 2023) using Firebase, a web application platform, were released as open source and used in motor learning research (Avraham et al., 2021; Tsay et al., 2022; Zhu et al., 2023). Remote experiments using web applications are an effective tool for bridging the gap between experiments using robotic manipulandum in the laboratory and general motor tasks outside the laboratory.

The influence of interventions that increase error or variability in remote motor tasks on motor learning has not been determined. Therefore, in this study, we aimed to determine the effects of interventions that visually increase errors and variability in remote experiments using web applications.

## 2 MATERIAL AND METHODS

### 2.1 Participants

People who understood Japanese and had laptops for web-based experiments were recruited into the study. A total of 48 people (30 men and 18 women; 25.5 ± 5.4 years old) participated in the study. Participants provided written informed consent via email to the authors before the commencement of the study. The experimental procedure used in this study was approved by the ethics committee of the institution with which the authors are affiliated. Participants completed the experiment, submitted the required files via email, and received a reward. Note that the number of participants who correctly completed the entire task was less than 48 because several quit the experiment within nine days, and others misunderstood the experiment procedure.

### 2.2 Motor tasks and protocols

The participants were instructed to use their laptops with trackpads and access the webpage for the experiments (Fig.1A). The motor tasks were developed by modifying an open-source package for online experiments referred to as “OnPoint” (Tsay et al., 2020). The motor task programs were edited in JavaScript and published via Firebase as a web application. The experimental protocol is shown in Fig.1B. Three motor tasks were conducted, and each task had a duration of three days (nine days in total). For each task, no intervention was provided on Day 1 and intervention to augment motor errors or variability was provided on Day 2 and 3. For each day, the procedure consisted of the first five trials of familiarization and the following 120 training trials. In the familiarization trials, the start and target positions differed from those in the training trials, to prevent the participants from learning the tasks. The final 20 trials of training trials on Day 3 were the test trials in which no intervention was provided. The number of trials was determined as follows: Participants could concentrate on completing the training (appropriately 10-15 minutes per day). The height *h* and width *w* of the screen were recorded and used as normalizing parameters, such as the target position on each day. In the JavaScript display coordinate system, the *x* and *y* positions of a point are the distances from the left and top of the window (Fig.1C).

**Figure 1.**
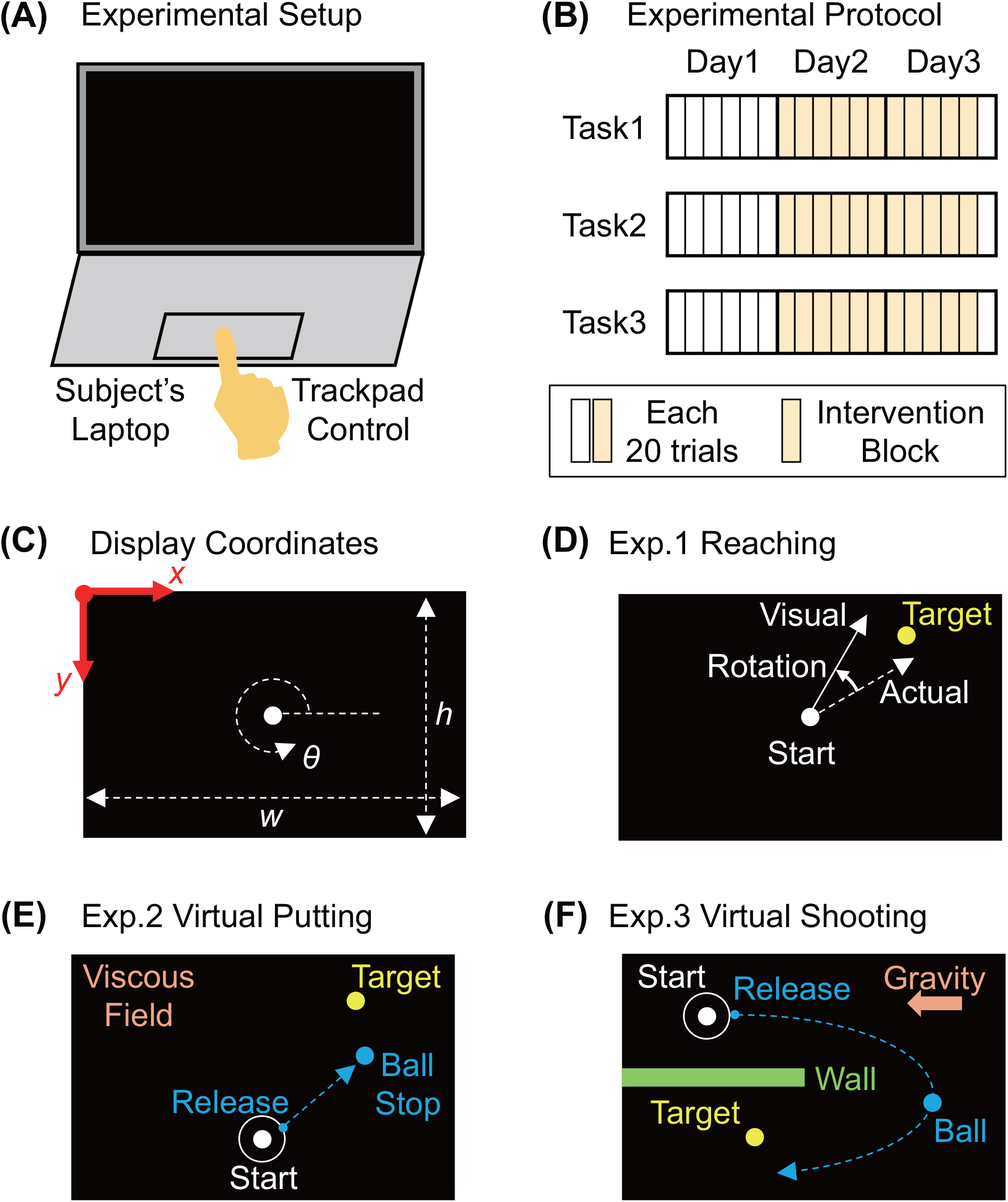
(A) Experimental setup. (B) Experimental protocol. (C) Display coordinates. (D)(E)(F) Schematic descriptions of tasks in Experiments 1, 2, and 3, respectively.

In Experiment 1, a reaching task with visuomotor rotation was conducted (Fig.1D). The start reaching position was 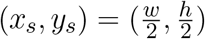. The distance from the starting position to the target was 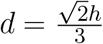. The target is circular, and its position was (*x*_*t*_, *y*_*t*_) = (*x*_*s*_ + *d* cos *θ*_*t*_, *y*_*s*_ − *d* sin *θ*_*t*_). In this study, *θ*_*t*_ was set to 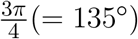 in the familiarization trials and to 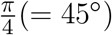 in the training trials for right-handed participants. For left-handed participants, *θ*_*t*_ was set to 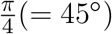 in the familiarization trials and to 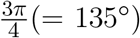 in the training trials. From participants’ viewpoint, 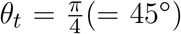 and 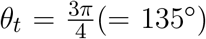 indicated the top-right and top-left directions from the starting position in the display, respectively. The radii of the cursor and target were 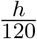 and 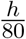, respectively. In the training trials, a constant visuomotor rotation was applied, where the visual hand movement angle was rotated at 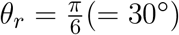 from the actual hand movement *θ*_*h*_.

In Experiment 2, a virtual curling or putting task was performed in which a ball was thrown using the cursor movement (Fig.1E). The starting position of curling was 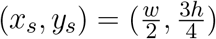. The distance from the starting position to the target was 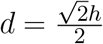. The target was circular, and its position was (*x*_*t*_, *y*_*t*_) = (*x*_*s*_ + *d* cos *θ*_*t*_, *y*_*s*_ − *d* sin *θ*_*t*_). Here, *θ*_*t*_ was set to 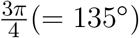 in the familiarization trials and to 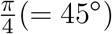 in the training trials for the right-handed participants. For the left-handed participants, *θ*_*t*_ was set to 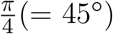 in the familiarization trials and to 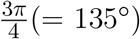 in the training trials. The radii of the cursor and target are 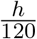 and 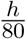, respectively. The cursor was controlled in the field circle of which the radius is 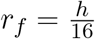, and the ball was released when the cursor reached the edge of the field circle. The release parameters of the ball such as the release speed and angle were defined depending on the cursor movement velocity at the time of release. The force acting on the ball after the release was a viscous force in the direction opposite to the motion of the ball.

In Experiment 3, a virtual ball-throwing task was conducted, similar to basketball shooting, in which a ball was thrown via the cursor movement (Fig.1F). The starting position of the shooting was 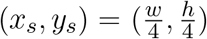 for the right-handed participants and 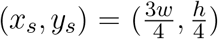 for the left-handed participants. The target shape was a line and its position was 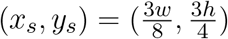 for the right-handed participants and 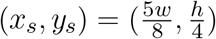 for the left-handed participants. The target length was 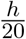 in the vertical direction, which was four times the radius of the ball. The cursor was controlled in the field circle, which had a radius of 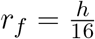, and the ball was released when the cursor reached the edge of the field circle. The forces acting on the ball after release were a viscous force in the direction opposite to the motion of the ball, and a constant gravitational force in the right-to-left direction for the right-handed participants and left-to-right for the left-handed participants. A wall was added to force the participants to avoid straight-line ball trajectories and use high-arch trajectories.

### 2.3 Intervention

In Experiment 1, the participants were divided into four groups: control (C), deterministic error augmentation (DEA), stochastic error augmentation (SEA), and bidirectional noise (BDN). For the DEA group, the actual reach angle error was multiplied by the constant *A* = 2. Thus, the visual angle from the starting position of the cursor position which was visually presented to the participant was *θ*_*v*_ = *θ*_*t*_ + *A*(*θ*_*r*_ − *θ*_*t*_). For the SEA group, the actual reach angle error was multiplied by a constant *A* plus random noise *ξ* from a uniform distribution [− (*A* − 1), (*A* − 1)] (Hasson et al., 2016). Thus, the visual angle from the starting position of the cursor position which was visually presented to the participant was *θ*_*v*_ = *θ*_*t*_ + (*A* + *ξ*)(*θ*_*r*_ − *θ*_*t*_). For the BDN group, the actual reach angle was added by random noise *ξ* from uniform distribution [−*B, B*], where 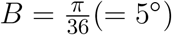. Thus, the visual angle from the starting cursor position which was visually presented to the participant was *θ*_*v*_ = *θ*_*h*_ + *θ*_*r*_ + *ξ*.

For Experiment 2, the participants were divided into four groups: control (C), speed noise (SN), angle noise (AN), and both noise (BN). For the SN group, the putting distance error was stochastically augmented by a constant *A* = 2 plus random noise *ξ* from the uniform distribution [− (*A* − 1), (*A* − 1)]. Thus, the visual putting distance from the release position was *d*_*v*_ = *d*_*r*_ + (*A* + *ξ*)*d*_*ae*_, where *d*_*r*_ was the distance from the release to the target and *d*_*ae*_ was the distance of the actual error. For the AN group, the actual release angle was manipulated in the same manner as in the SEA group in experiment 1. In particular, the visual angle from the release position was *θ*_*v*_ = *θ*_*opt*_ + (*A* + *ξ*)*θ*_*ae*_, where *θ*_*opt*_ was the optimal release angle and *θ*_*ae*_ was the angle of actual error. For the BN group, the error in putting distance and release angle were manipulated as described above for the SN and AN group.

For Experiment 3, the participants were divided into three groups: control (C), speed noise (SN), and angle noise (AN) group. Two noise levels (high and low) were provided to the SN and AN groups. The SN and AN groups were further subdivided into speed noise high (SNH), speed noise low (SNL), angle noise high (ANH), and angle noise low (ANL) groups. All noises were drawn randomly from the normal distribution with a mean of 0 and standard deviation of *σ*. For the AN group, the standard deviation was as follows: 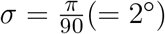 and 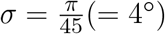 for low and high noise, respectively. In the SN group, the standard deviation was set to *σ* = 0.05*v*_min_ and *σ* = 0.1*v*_min_ for low and high noise, respectively, where *v*_min_ is the minimum release speed required to hit the target (Brancazio, 1981), which can be calculated using the equation of motion of the ball flight and release position.

### 2.4 Data collection and analysis

The height *h* and width *w* of the screen were recorded on each day of the three experiments. In Experiment 1, the actual reaching angle and visual reaching angle, which was manipulated with constant visuomotor rotation and an intervention depending on the groups, were recorded. In Experiment 2 and 3, ball release parameters and visual ball arrival positions were recorded, which are related to each other via an equation of motion for ball movements. In Experiment 3, the solution manifold (Nakano et al., 2020) in the space of release speed and angle was drawn with the measured data to determine the release strategies of throwing. All data analysis was performed using MATLAB.

### 2.5 Statistics

Statistical analysis was performed on motor performance in the final 20 trials (i.e., the test phase) of each experiment using a two-sample t-test. The calculations were performed using the MATLAB function “ttest2”.

## 3 RESULTS

### Experiment 1

In Experiment 1, the mean reaching angle error for each group was calculated for each block (consisting of 20 trials), as shown in Fig.2. As a constant visuomotor rotation was applied from Block 2, the angle error increased from Blocks 1 to 2 and rapidly decreased from Block 2. Note that the angle error in the SEA group decreased more gradually than in the other groups. The interventions were applied from the start of Day 2; however, the angle error of the DEA, SEA, and BDN groups did not differ significantly from that of the constant group. The statistical analysis revealed no significant differences between any two of the four groups (C, DEA, SEA, and BDN) in the test phase on Day 3.

**Figure 2.**
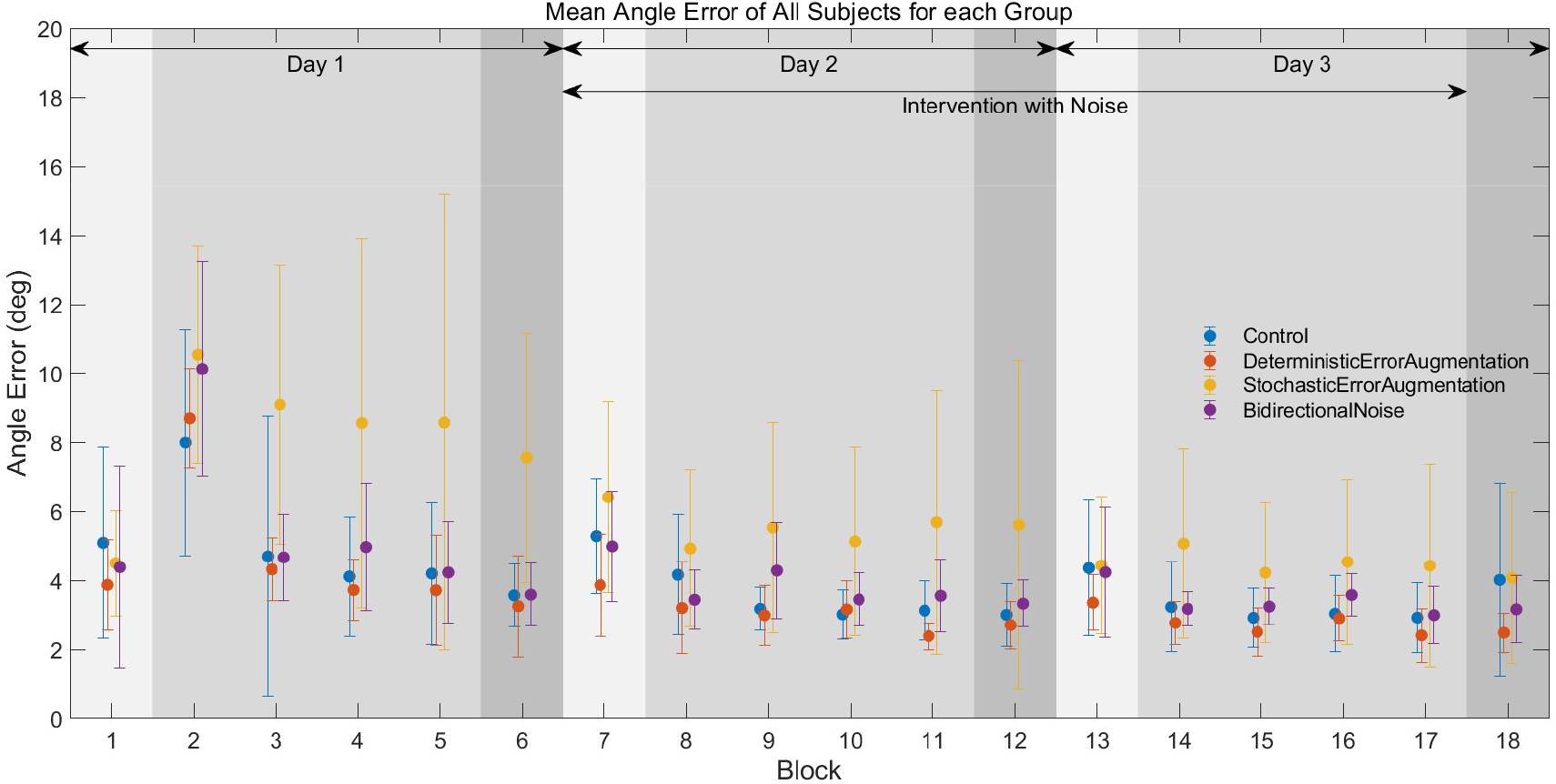
Mean reaching angle error for each group calculated for each block (20 trials). The horizontal axis indicates the number of blocks in the learning process and the vertical axis indicates the angle error of reaching.

### Experiment 2

In Experiment 2, the mean performance error in putting angle and distance for each group was calculated for each block (consisting of 20 trials), as shown in Figs.3 and 4, respectively. Fig.3 revealed that the angle errors of the AN and BN groups were larger than those of the C and SN groups in the Block 7-17 owing to the intervention. However, the statistical analysis revealed no significant differences in angle error between any two of the four groups (C, AN, SN, and BN) in the test phase on Day 3. Fig.4 revealed that the distance errors of the SN and BN groups were larger than those of the C and AN groups in Block 7-17 owing to the intervention. However, statistical analysis revealed no significant differences in distance error between any two of the four groups (C, AN, SN, and BN) in the test phase on Day 3.

**Figure 3.**
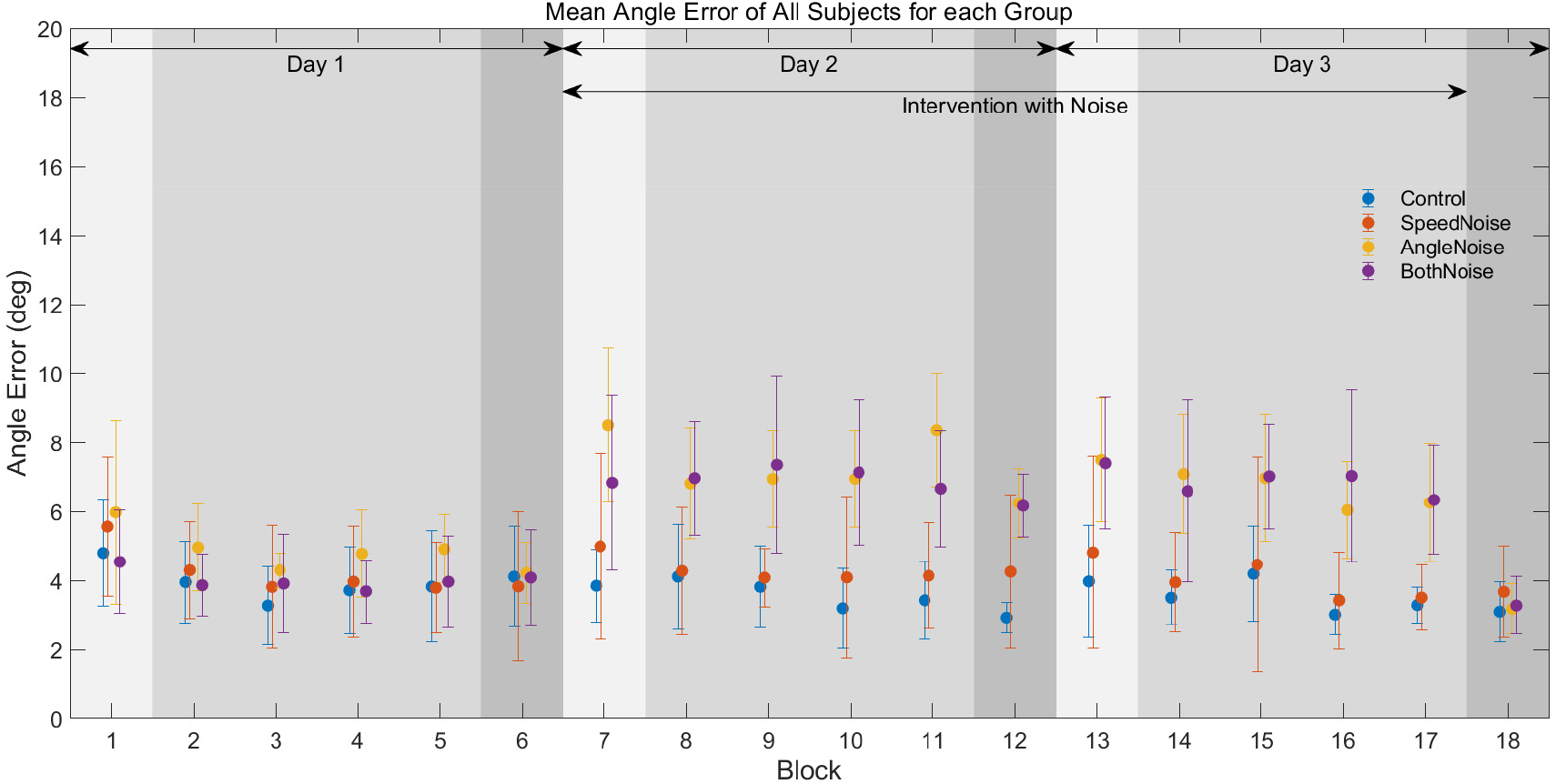
Mean putting angle error for each group calculated for each block (20 trials). Experiment 3

**Figure 4.**
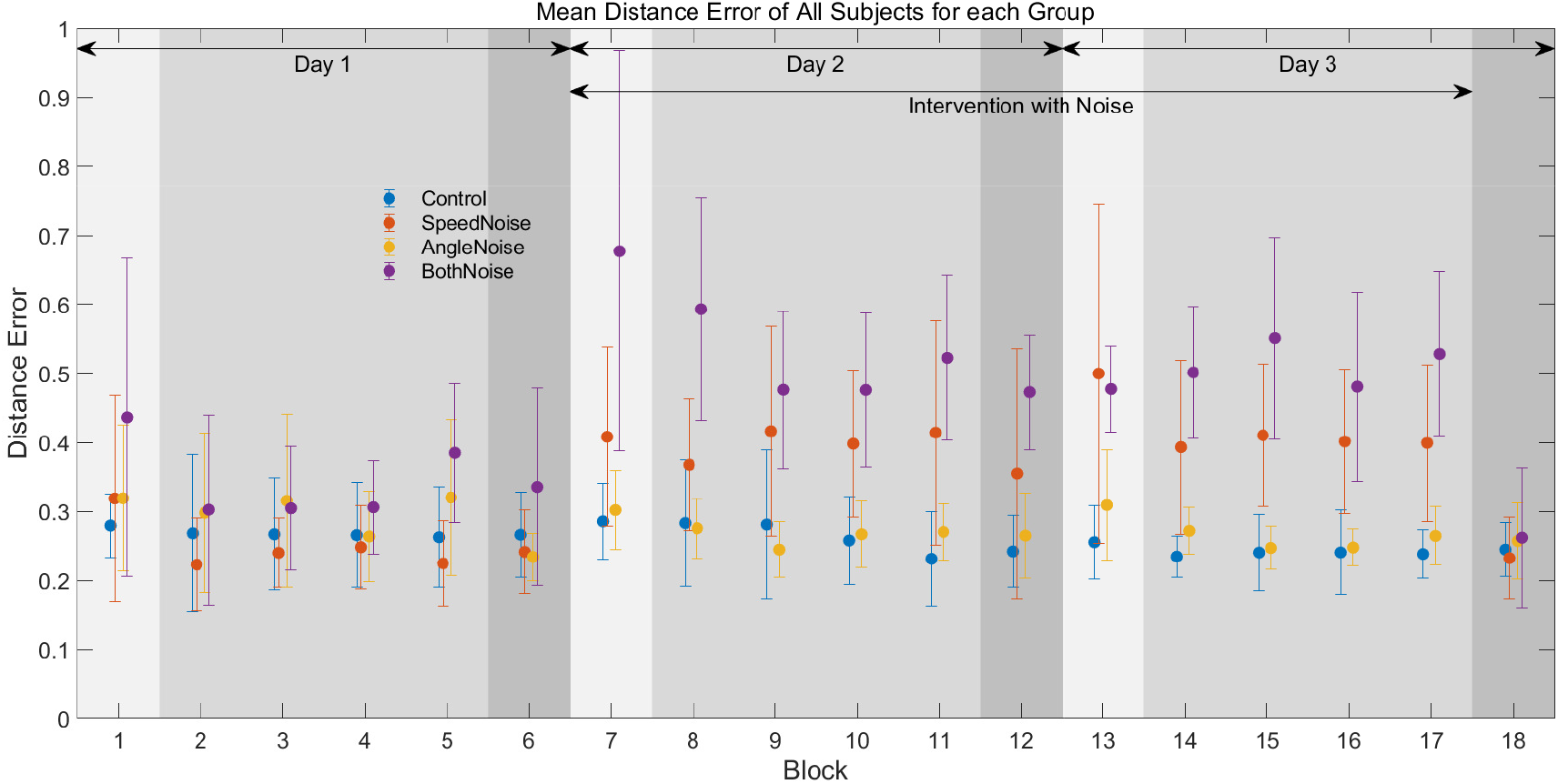
Mean Putting distance error for each group calculated for each block (20 trials).

In Experiment 3, the solution manifold and actual release parameters for pre- and post-intervention blocks are shown in Fig.5. The colormap indicates the distance error from the target and the white area indicates zero-error from the target. Noted that the mean values of the parameters for all participants with respect to the display size, such as the target positions, start positions, and target sizes were used to draw the solution manifold, such that all the participant data could be visualized within a single figure. Thus, the solution manifold (Fig.5) does not completely correspond to that calculated daily for each participant. The shape of the solution manifold revealed that the optimal release strategy was uniquely determined when the release angle was determined; therefore, the release angle for each group was quantitatively examined. The mean shooting angle for each group was calculated for each block (20 trials), as shown in Fig.6. However, the statistical analysis revealed no significant differences in the release angle between any two of the four groups (C, SN, and AN) in the test phase on Day 3.

**Figure 5.**
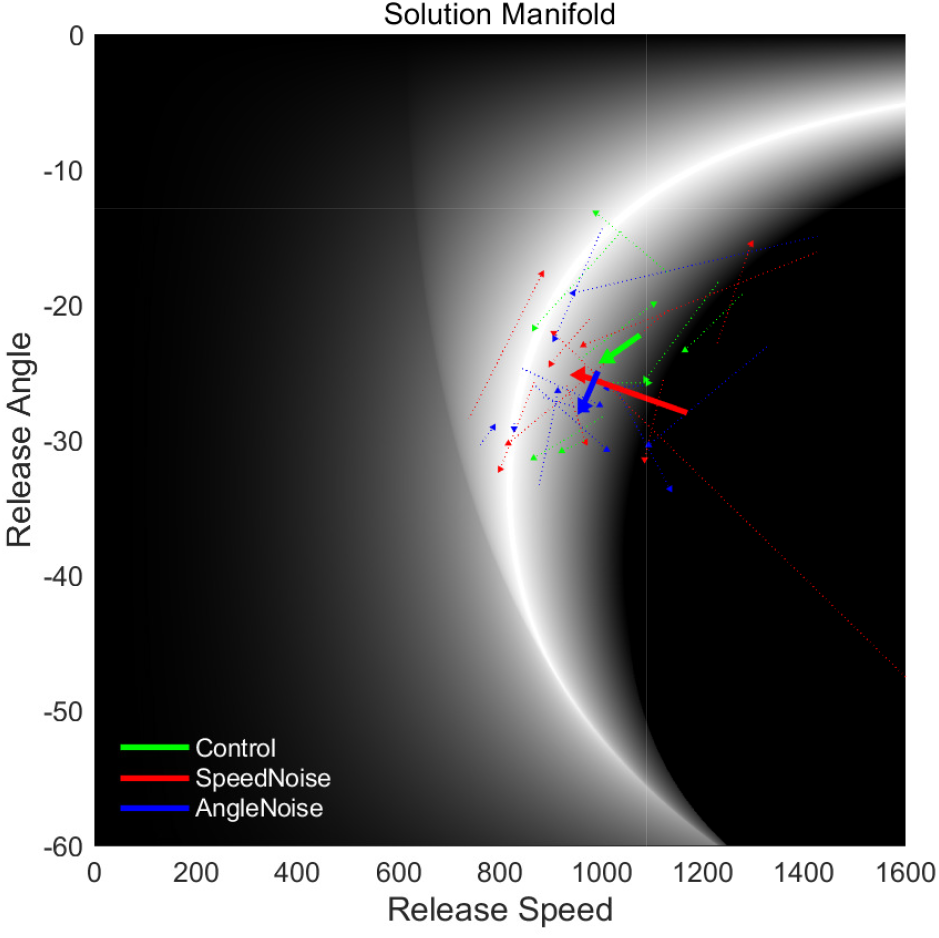
Solution manifold and the actual release parameters for the pre- and post-intervention block. The colormap indicates the distance error from the target, and the white area indicates the zero-error from the target. The arrows indicate the change from the test phase on Day 1 (Block 6) to the test phase on Day 3 (Block 18). The solid lines indicate the mean between all participants in each group, and the dotted lines indicate each participant in each group. The green, red, and blue lines indicate the group of control, speed noise, and angle noise, respectively.

**Figure 6.**
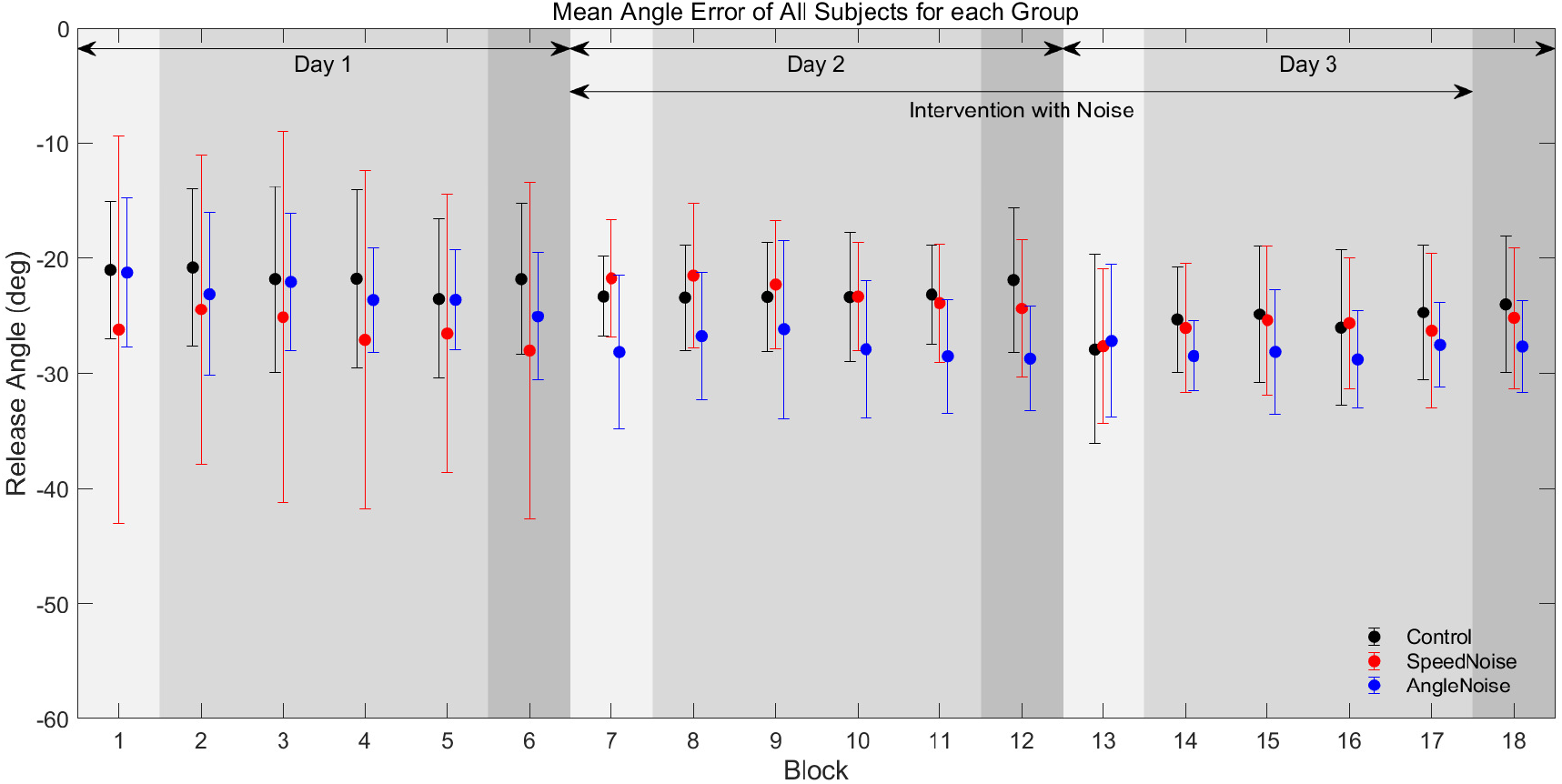
Mean shooting angle for each group calculated for each block (20 trials).

## 4 DISCUSSION

In this study, we aimed to determine the effects of interventions that visually increase errors and variability in remote experiments using web applications. None of the experiments revealed the positive effects that are often reported for laboratory tasks.

### ERROR AUGMENTATION FOR IMPROVING MOTOR ACCURACY

We hypothesized that (1) error amplification (deterministic and stochastic) improves motor accuracy, (2) random noise in both directions relative to the error does not improve motor accuracy, and (3) intervention on various variables (e.g., speed and angle) improves the accuracy of the corresponding variables. However, the results of Experiments 1 and 2 disprove these hypotheses. These negative effects are consistent with those reported in the motor learning studies on rowing simulators (Gerig et al., 2019). In the abovementioned study, the significant negative effects were attributed to large inter-subject variability during practice, small differences between the control and intervention conditions, and lack of error-based learning in the dominant mechanism for reproducing oar trajectory. Experiment 1 in this study exhibited no differences between conditions during training in the intervention conditions (Fig.2). This is because of the large inter-subject variability, even within the conditions. Conversely, in Experiment 2, the differences between the conditions were clear during training in the intervention conditions (Figs.3 and 4) owing to a small inter-subject variability within the conditions. The intervention negatively influenced motor learning irrespective of the amount of inter-subject variability, whereas other factors produced negative effects in this study. The intervention and on-screen behavior in the previous study (Hasson et al., 2016), which used visual error amplification intervention in a virtual throwing task, was almost identical to that used in the present study; however, the ball was thrown at elbow flexion using a manipulandum. Thus, the haptic and proprioceptive information obtained during experiments in the previous study was significantly different from that used in the present study. Haptic error amplification has reported to have a more positive impact on motor learning than that of visual error amplification (Marchal-Crespo et al., 2019), and it is highly probable that the link between motor performance and haptic and/or proprioceptive information is critical for learning.

### VARIABILITY AUGMENTATION FOR GUIDING MOTOR STRATEGY

We hypothesized that intervention with noise, which increases variability, would lead to solutions that are less influenced by noise (i.e., a more noise-tolerant strategy). However, the results of Experiment 3 disagreed with this hypothesis. This result is inconsistent with previous research showing that random noise, which implicitly increases motor variability, drives motor learning toward decreasing motor variability (Mehler et al., 2017; Thorp et al., 2017). Given that the noise magnitude is directly dependent on the hand postures (i.e., angle) in these studies, the relationship between motor performance (task-space) and hand posture (joint space) may be more readily observed via proprioceptive sensing. In this study, on the other hand, the magnitude of the noise does not depend directly on the hand motions, and the effect of the noise on motor performance is determined by the shape of the solution set (i.e., the mapping from release space to task space). The shape of the solution manifold (Fig.5) indicates that the minimum-speed strategy (i.e., −40 ∼ −30°) minimize the effect of angle error, and using a high-arch strategy (i.e., −10 ∼ 0°) minimize the effect of speed error (Nakano et al., 2020). Although those who had learned sufficiently, such as experienced basketball players throwing free throws (Nakano et al., 2020) or the general public who learned virtual skittle throwing for several days (Sternad et al., 2011; Zhang et al., 2018), were able to determine a suitable release strategy for the solution manifold; the learning time in this experiment may have been insufficient to find a suitable strategy for artificially adding variability. Therefore, it is suggested that the effect of the intervention on training is more positive in situations where the relationship between the variable being intervened (body space) and the motor performance (task space) is direct.

### LIMITATION

A limitation common to all the experiments in this study was the accuracy of measuring cursor kinematics. Although specialized systems designed for motion measurement (e.g., optical motion capture systems and robotic manipulandums) exhibit very high temporal and spatial resolutions, common laptop trackpads are less accurate than these systems in terms of kinematics. Therefore, the intervention to increase motor errors, where the task programs identify and manipulate the error from the measured motion kinematics, may not have been sufficiently accurate. Another limitation common to all experiments was the motivation and concentration of the participants. When participating in an experiment in a new location or with an experimental apparatus while being observed by strangers, most participants were more nervous and focused than under the present experimental conditions. In this study, the participants were instructed to maximally focus on the task. However, the degree of execution may have differed from participant to participant. To clarify whether these two limitations are the main causes of the negative results, further research is required to verify whether the same results are obtained if a task almost identical to the present task is performed using a laboratory robot manipulandum.

## AUTHOR CONTRIBUTIONS

NN: Conceptualization, Data curation, Formal analysis, Funding acquisition, Investigation, Methodology, Resources, Software, Validation, Visualization, Writing – original draft, Writing – review & editing; AM: Funding acquisition, Project administration, Supervision, Writing – review & editing.

## DATA AVAILABILITY STATEMENT

The datasets analyzed for this study can be found in the figshare: https://figshare.com/articles/dataset/Nakano2024Data_zip/25795417.

